# A Humanized Antibody against LRG1 that Inhibits Angiogenesis and Reduces Retinal Vascular Leakage

**DOI:** 10.1101/2020.07.25.218149

**Authors:** David Kallenberg, Vineeta Tripathi, Faiza Javaid, Camilla Pilotti, Jestin George, Sterenn Davis, Jack WD Blackburn, Marie O’Connor, Laura Dowsett, Chantelle E Bowers, Sidath Liyanage, Morgane Gourlaouen, Alexandra Hoeh, Filipa Mota, David Selwood, James Bainbridge, Adnan Tufail, Vijay Chudasama, John Greenwood, Stephen E Moss

**Author notes:** DK and VT equal first authors.

## Abstract

Pathological angiogenesis contributes to morbidity in a number of diseases including cancer, diabetic retinopathy and the neovascular form of age-related macular degeneration, leading to significant efforts to develop effective anti-angiogenic therapeutics for these conditions. The field is dominated by inhibitors of vascular endothelial growth factor (VEGF), yet angiogenesis can also be driven and modified by other factors. We have previously demonstrated that leucine-rich alpha-2-glycoprotein 1 (LRG1) contributes to abnormal vessel growth by activating a TGFß switch. Here we report the development and characterisation of a function-blocking fully humanised IgG4 and its Fab fragment, that bind to LRG1 with high affinity and specificity and inhibit vascular leakage in the mouse model of laser-induced choroidal neovascularisation. In summary, we have developed a therapeutic antibody that targets a VEGF-independent signalling axis, which may be effective in a number of conditions either as monotherapy or in combination with other vascular targeted therapies.

## INTRODUCTION

The neovascular form of age-related macular degeneration (nvAMD) and diabetic eye disease are among the most prevalent forms of sight loss globally, with both having a significant impact on quality of life and presenting major health care challenges. Despite their distinct etiologies, both conditions are characterized by pathological changes to the ocular vasculature, which in turn has driven the search for and development of anti-angiogenic therapeutics.^1–3^ In nvAMD the blood vessels of the choroid typically penetrate Bruch’s membrane and the retinal pigment epithelium, from where they may anastomose with the retinal capillary network to create a disorganized and dysfunctional vascular tangle. The edema that typically accompanies these changes causes swelling and often irreparable damage to the macula. In diabetic eye disease the vascular defects tend to be confined to the retinal vessels where pericyte dropout, microaneurysms, neovascular tufts, edema and ultimately neo-angiogenesis may also cause irreversible vision loss. In both conditions the fragile neovessels may also hemorrhage with devastating and potentially irreversible effects on vision.

In both conditions one of the major drivers of pathology is vascular endothelial growth factor (VEGF), hence anti-VEGFs have become the go-to therapeutic option for most clinicians.^4,5^ In nvAMD approximately 85% of patients derive at least some benefit from monthly or bi-monthly intravitreal injections of VEGF-blocking therapeutics such as aflibercept, ranibizumab and bevacizumab. Favorable outcomes are less frequent in diabetic eye disease where around 50% of patients fail to respond to these interventions. So while the anti-VEGFs have been a clear success there remains a significant unmet clinical need, along with the implication that VEGF-independent mechanisms must be playing a role in development of the vascular pathology in many patients. Certainly, other factors such as the angiopoietins, PDGF, TGFß and the FGFs have all attracted the attention of drug developers, and a number of blocking antibodies and bi-specifics have been, and continue to be, developed and tested in clinical trials. Among these TGFβ presents particular challenges, not least because of the context dependency of its activities and its multiple housekeeping roles in different ocular tissues.^6,7^

In previous work we identified a novel regulator of TGFβ signaling named LRG1, that by altering the stoichiometry of the TGFβ receptor complex in vascular endothelial cells drives a switch in signaling that contributes to pathogenic angiogenesis and vessel destabilisation.^8^ LRG1 is a secreted glycoprotein synthesised primarily by the liver under normal conditions, but which is frequently induced and expressed at sites of pathological angiogenesis. Consistent with this, we observed that LRG1 levels are significantly elevated in the vitreous of patients with diabetic retinopathy,^8^ it is present in both the aqueous and vitreous proteome in wet AMD^9^,^10^ and there are numerous reports of increased serum LRG1 in solid cancers,^11–13^ suggesting that it may also contribute to tumour angiogenesis. LRG1 is dispensable for developmental angiogenesis as null mutant mice develop a normal vasculature, are fertile and have a typical lifespan. In contrast, we demonstrated that pathological angiogenesis is significantly reduced in the absence of LRG1 in a number of inducible disease models. These observations suggested that LRG1 is a promising therapeutic target in pathological angiogenesis, particularly as LRG1 blockade inhibits the pro-angiogenic activity of TGFβ, without interfering with its many homeostatic roles, and may therefore address the challenge of treating anti-VEGF refractive patients.

## MATERIALS AND METHODS

### Antibody generation and humanisation

Monoclonal antibodies (mAbs) against recombinant human LRG1 were generated in mice using conventional hybridoma technology (EMBL, Italy). More than 100 mAbs were isolated and taken forward for analysis by SPR and in functional assays. The humanisation and deimmunisation of lead monoclonal antibody 15C4 were performed by Abzena (Cambridge, UK). Recombinant human, cynomolgus and mouse LRG1 were expressed as His-tagged fusions and purified by FPLC from the culture supernatants of stable-transfected CHO producer cell lines.

### Surface plasmon resonance

The binding kinetics and affinities of 15C4, Magacizumab and MagaFab for recombinant human, cynomolgus or mouse LRG1 were determined by SPR (Biacore T200, GE Healthcare). All samples were buffer exchanged into running buffer HBS-EP+, pH 7.4 (GE Healthcare). The respective LRG1 proteins were immobilised directly onto a CM5 biosensor chip by amide coupling at pH 5.0. Targeting a low immobilisation level of ~100 RU to avoid avidity. Antibodies were injected over immobilised LRG1 for 120 seconds at 30μL/min at increasing concentrations using a single-cycle kinetics methodology, with an extended dissociation time of 3600 seconds. A 1:1 fit model was applied to determine the binding kinetics (k_on_ and k_off_), binding affinities were obtained using the formula: K_D_ = k_off_/k_on_.

### Co-culture angiogenesis assay

The angiogenesis assay was performed according to manufacturer’s instructions (Cellworks). Briefly, human umbilical vein endothelial cells (HUVECs) and human fibroblasts were seeded, and the plate was incubated at 37 °C in 5% CO_2_ overnight. Cells were treated with 15C4 (667 nM), and control IgG (667 nM) diluted in growth medium. Medium was replaced every two days for 11-14 days of culture, during which time the endothelial cells reorganise to form tubules that resemble microvessels formed during angiogenesis. Cells were washed with PBS and fixed with ice cold 70% ethanol for 30 min at room temperature, then immunostained with mouse anti-human CD31 (Cellworks). Cells were imaged using a TiE Nikon Inverted Fluorescence Microscope. Angiogenesis was quantitated using Angiosys (TCS Cellworks). The total tubule length for the treated samples was normalized to equivalent controls. Three independent experiments were carried out with a mean of 12 wells being analysed for each treatment. One-way ANOVA was used to determine the statistical significance between groups.

### Metatarsal assay

Metatarsal bones were isolated from mouse embryos at day 17 – 18 and dissection was performed in sterile-filtered 10% heat inactivated foetal bovine serum (FBS) (Gibco) in PBS. 24-well plates were coated with sterile-filtered 0.1% gelatine (Sigma, cat. no. G1393) diluted in 1 × PBS. Each bone was plated in the middle of the well with α-MEM medium (MEM α Med 1 + w/Earle’s Salts Glut Nucleosides, Invitrogen, cat. no. 22571020) supplemented with 10% FBS and 100 U/mL Penicillin and 100 mg/mL Streptomycin (Gibco) and incubated overnight at 37 °C in 5% CO_2_. The metatarsals were left undisturbed for 3 days allowing them to adhere to the wells. On day 3, 5, 7 and 9 metatarsals were treated with antibodies as detailed in the figure legends, diluted in α-MEM medium. On day 11 the conditioned medium was collected and stored at −80 °C for future analysis. Bones were fixed with 4% paraformaldehyde (PFA) in PBS for 30 min at room temperature. Non-specific binding of antibodies was inhibited by addition of blocking buffer (10% BSA and 0.1% Triton X-100 (Sigma) in PBS) for 1 h at room temperature. This was replaced with primary antibody (rat anti-mouse CD31, 1:100) diluted in 5% BSA and 0.1% Triton in PBS and left overnight at 4 °C. Excess antibody was washed off with 0.1% Tween-20 in PBS and then replaced with 5% BSA and 0.1% Triton in PBS containing the secondary antibody (AlexaFluor 488 Goat anti-rat IgG, 1:1000) for 2 h at room temperature. Four washes were done with 0.1% Tween in PBS for 1 h. Bones were imaged using a TiE Nikon Inverted Fluorescence Microscope. Angiogenesis was quantified using Angiosys software (TCS Cellworks). The number of vessel branches and the total tubule length for the treated samples was normalized to equivalent controls. Metatarsals from three independent litters were used and one-way ANOVA with multiple comparisons was performed to determine statistical significance between test groups.

### Fab preparation

Magacizumab (75.4 mg/mL) in digestion buffer was reacted with a 1/10 amount (wt/wt) of immobilised papain (250 μg/mL of gel) by incubation for 24 h at 37°C whilst shaking (1100 rpm), in a buffer containing 20 mM sodium phosphate monobasic, 10 mM disodium EDTA and 80 mM cysteine·HCl (pH 7.2). The cysteine·HCl was incorporated immediately before Magacizumab digestion. After digestion, the resin was separated from the digest using a filter column and washed with PBS (pH 7.0) three times. The digest was combined with the washes and the buffer was exchanged completely for PBS (pH 7.4) using diafiltration columns (10,000 Da MWCO) and the volume adjusted to 2 mL. The sample was then applied to a NAb protein A column (Thermo Scientific). The Fab fraction (MagaFab) was eluted according to manufacturers’ protocol, the column washed three times with PBS (pH 7.4) and the Fc fraction, eluted four times with 0.2 M glycine·HCl, pH 2.5, which was neutralised with 10% of the volume of a 1 M Tris, pH 8.5 solution. The MagaFab fraction was combined with the washes, and both Fab and Fc solutions were buffer exchanged into PBS using diafiltration columns (10,000 Da MWCO). The digests were analysed by SDS-PAGE and LCMS to reveal formation of MagaFab; observed mass 47,461 Da.

### Preparation of MagaFab conjugates

Full details of the MagaFab-Cy5.5 preparation are provided in the **supplementary text**. Briefly, to a solution of MagaFab (20 μM, 200 μL) in BBS (25 mM sodium borate, 25 mM NaCl, 0.5 mM EDTA, pH 8.0) was added TCEP (20 μL, 20 mM in dH2O, 100 eq.), and the reaction mixture incubated for 4 h at 37°C under mild agitation (300 rpm). After this time excess TCEP was removed by diafiltration into fresh BBS buffer using PD Minitrap G-25 columns (GE Healthcare) and the concentration was corrected to 20 μM. Pyridazinedione^14^ (PD) (1 μL, 20 mM in DMSO, 5 eq.) was added to the solution of reduced MagaFab at 21°C and the solution incubated for 16 h. Excess reagents were removed by repeated diafiltration into fresh PBS buffer (140 mM sodium chloride and 12 mM sodium phosphates at pH 7.4) using VivaSpin sample concentrators (GE Healthcare, 10,000 Da MWCO), and the volume was corrected to 200 μL. The samples were analysed by SDS-PAGE gel and UV-Vis spectroscopy, which was used to determine a Pyridazinedione to antibody ratio of 0.99. Following this, Sulfo-Cyanine5.5 Azide (Lumiprobe) (1 μL, 20 mM in DMSO, 5 eq.) was added and the reaction mixture was incubated at 21°C for 4 h. The excess unbound fluorophore was removed by repeated diafiltration into fresh PBS using diafiltration columns (GE Healthcare, 10,000 Da MWCO). Following this, sample analysis by SDS-PAGE gel and UV-vis spectroscopy revealed conversion to the desired MagaFab-PD Sulfo-cyanine5.5 conjugate with a fluorophore to antibody ratio of 0.93.

### Determination of fluorophore to antibody ratio (FAR)

UV-vis spectra were recorded on a Varian Cary 100 Bio UV-visible spectrophotometer, operating at 20°C. Baseline correction was performed using sample buffer as a blank. Calculation of fluorophore to antibody ratio (FAR) follows the formula below, where with Ɛ_280_= 70000 M^−1^ cm^−1^ for Fab, Ɛ_691_= 235000 M^−1^ cm^−1^ for Sulfocyanine5.5 and 0.11 as a correction factor for the dye absorption at 280 nm.

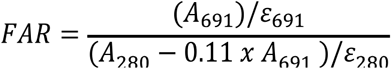

### Laser-induced choroidal neovascularisation

Laser-induced choroidal neovascularization (CNV) and quantification using fundus fluorescein angiography was performed as described previously.^8^ Each eye received three laser lesions at two to three disk diameters away from the optic nerve head with a slit-lamp-mounted diode laser system (Keeler microlase, Windsor, UK; wavelength: 810 nm; laser settings: 210 mW power, 100 ms duration, 100 μm spot diameter). Mice then received intravitreal injections of antibodies at doses indicated in the figure legends, immediately following the laser burn. Seven days after injury, mice were anaesthetized and examined by fundus fluorescein angiography (FFA). Images from the early phase (90 s after fluorescein injection) and late phase (7 min after injection) were obtained using a Micron III retinal imaging microscope with appropriate filters (Phoenix Research Labs, Pleasanton, USA). The pixel area of CNV-associated hyperfluorescence was quantified for each lesion using ImageJ version 1.44i image analysis software (NIH, Bethesda, MD). Images were blinded for analysis, and lesion perimeters were hand drawn to measure pixel area positive for fluorescein. One-way ANOVA with Tukey’s multiple comparison test was used to determine statistical significance between antibody treatment groups.

### Epitope mapping

Epitope mapping was performed by ProImmune (Oxford, UK) using a peptide microarray.

### RNAScope^®^ in situ hybridisation

FFPE retinal samples or formalin-fixed flat-mounted metatarsals were processed using the Multiplex Fluorescent Kit v2 (ACDBio), followed by TSA^®^ signal amplification (PerkinElmer). Slides were hybridised with probes specific to Mm*Lrg1* or Hs*LRG1*, and quality of signal and tissue determined using positive (Mm*Ppib*) and negative (*DapB*) probes, supplied by the manufacturer. In situ hybridisation was immediately followed by immunohistochemistry for antibody-mediated detection of vessel marker proteins CD31 and/or collagen IV in the same samples.

## RESULTS

### Functional blockade of LRG1

Using conventional hybridoma technology we first generated a panel of more than 100 monoclonal antibodies (mAbs) against recombinant glycosylated human LRG1 and assessed their binding kinetics by surface plasmon resonance (SPR), and their efficacy in blocking angiogenesis using an *in vitro* endothelial/fibroblast co-culture assay. Using selection criteria of equilibrium dissociation constant (K_D_) < 1 nM, and > 20% inhibition of endothelial tube formation respectively, we identified 12 antibodies that underwent in vivo functional testing using laser-induced choroidal neovascularisation (CNV) in mice (data not shown). Of these mAbs, a single antibody coded 15C4 emerged with the most favourable profile. 15C4 was purified from hybridoma supernatants by FPLC, and was observed, using single cycle kinetics on SPR, to have an association rate (k_on_) of 1.15 × 10^5^ M^−1^ s^−1^, a dissociation rate (k_off_) of 1 × 10^−5^ s^−1^ and a high K_D_ of 8.696 × 10^−11^ M, highlighting its potential as a therapeutic antibody (Table 1 and Supplementary text Figure S1). Additionally, SPR showed that 15C4 has high affinities for monkey and mouse LRG1, allowing functional evaluation of its activity in non-human species. In functional tests 15C4 significantly reduced tubule growth in the endothelial/fibroblast assay (**Figure 1a**) and exhibited comparable dose-dependent inhibition of vessel growth in the ex vivo mouse metatarsal angiogenesis model (**Figure 1b**). A combination of RNAScope^TM^ and immunofluorescence showed that *Lrg1* mRNA expression was readily detectable in cells closely associated with the sprouting vessels (Supplementary text Figure S2).

**Table 1.**
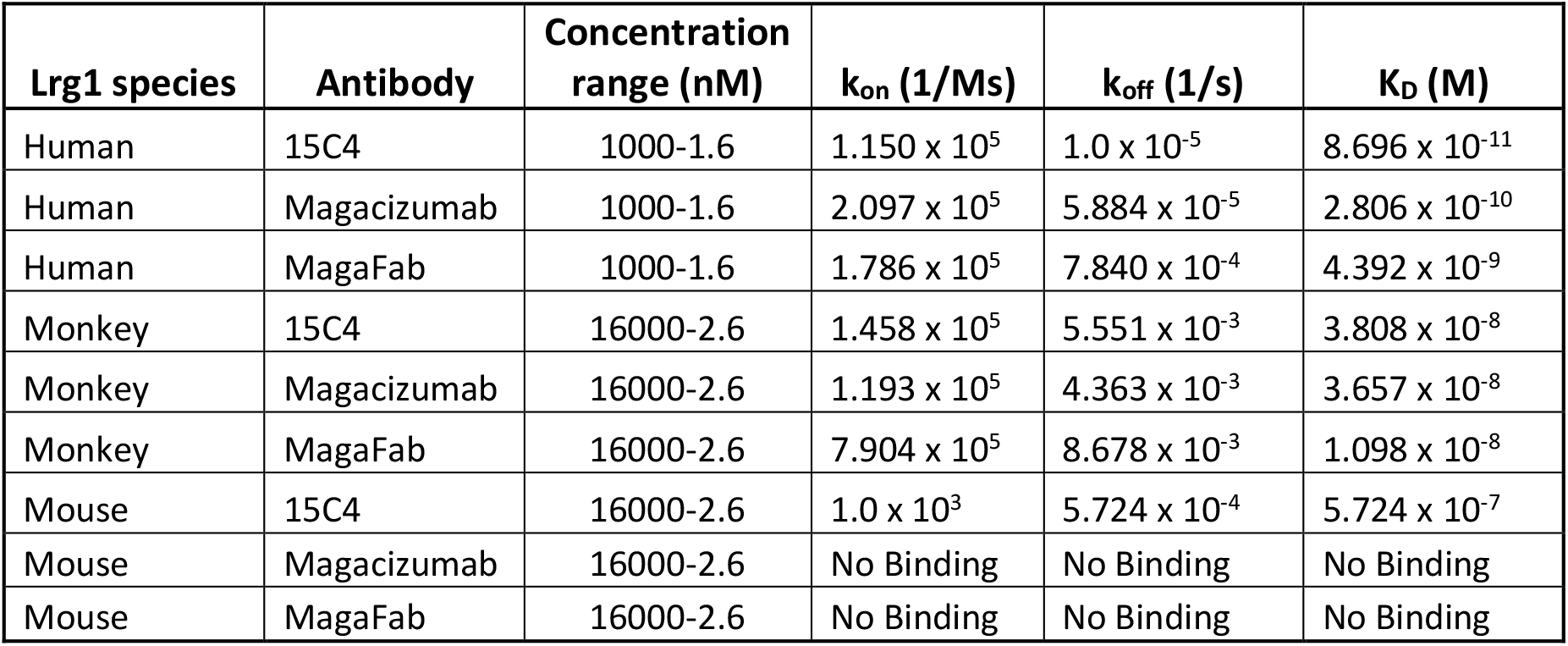
SPR binding kinetics of antibodies for recombinant LRG1 across species. The binding affinities of 15C4, Magacizumab and MagaFab for recombinant human, monkey or mouse LRG1 were determined by SPR (Biacore T200). Binding kinetics were determined using a one-to-one fit model.

**Figure 1.**
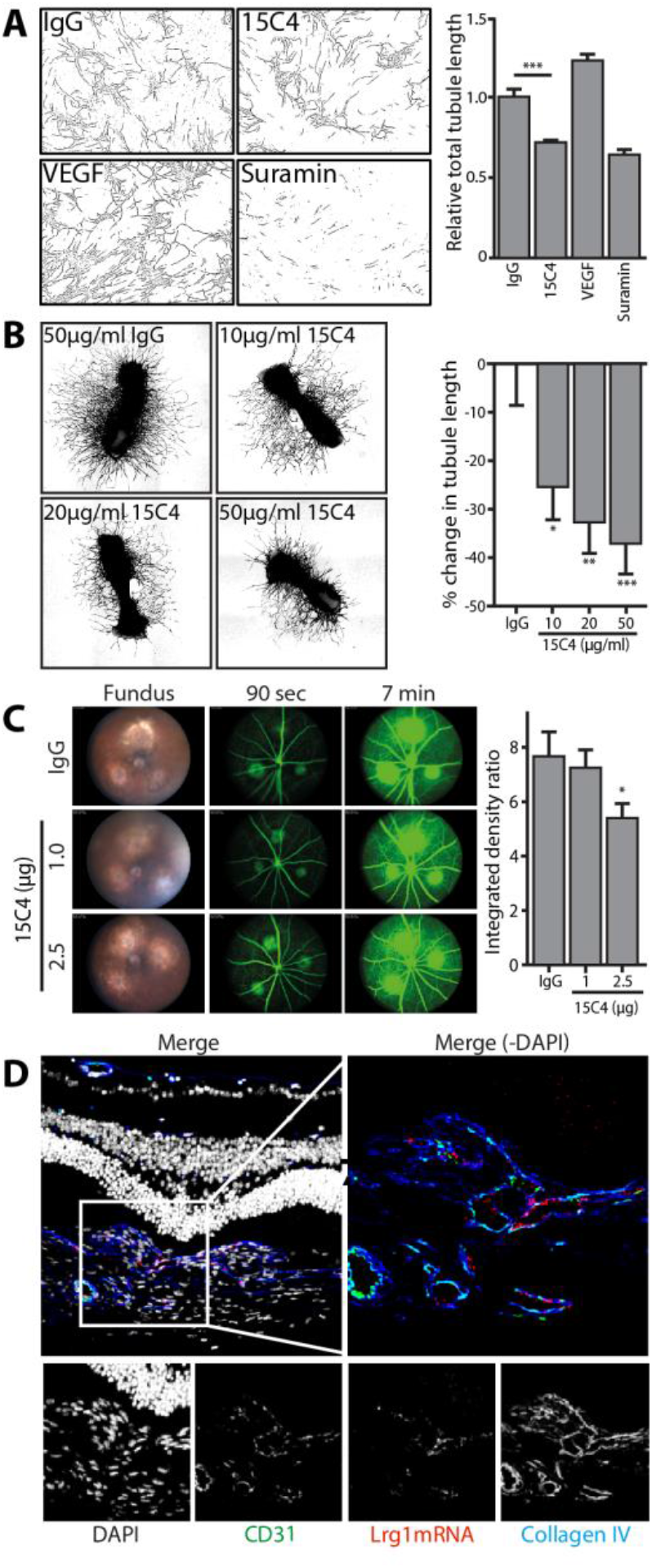
Inhibition of angiogenesis and vascular leakage with a monoclonal antibody against LRG1. (**a**) Co-cultured HUVECs and fibroblasts were treated with 15C4 or controls for 11 days, allowing for the formation of microtubules. Post-treatment, vessels were stained with anti-CD31 and analysed with AngioSys 2.0. A significant reduction in tubule length was observed with LRG1 blockade by 15C4 (100 μg/mL) compared to IgG controls (*\*\*\*P* < 0.0001, one-way ANOVA). (**b**) Metatarsals from wild type C57B6 mice were treated with doses of 15C4 as indicated with mouse IgG as a control, and immunostained for CD31 to reveal vessels. A significant reduction in total tubule length was observed in 15C4 treated metatarsals when compared to IgG (\**P* < 0.05, \*\**P* < 0.01, \*\*\**P*= 0.0023, one-way analysis of variance). (**c**) C57bl/6 mice with laser-induced choroidal neovascularisation were injected with 15C4 or IgG control immediately following laser burns. Fluorescein angiography was done on day 7. CNV leakage is significantly reduced in eyes treated with 2.5 μg 15C4 compared to IgG controls (one-way ANOVA followed by Tukey’s multiple comparison test, \**P* < 0.05). (**d**) Wild type mouse retinas were lasered and then after 7 days fixed and processed for analysis using a combination of RNAscope (to detect *Lrg1* mRNA) and immunocytochemistry (for CD31 and collagen IV). The merged image shows *Lrg1* mRNA expression in the endothelial cells within the lesion.

Having demonstrated the modulatory effects of 15C4 on vessel function in in vitro and explant models, we wanted to see if LRG1 blockade in vivo could impact on a primary defect associated with diseased vessels, namely leakage, as vasogenic oedema is a major pathological outcome in several forms of retinal vascular disease. Wet AMD is characterised by CNV and is associated with leakage of fluid and blood from abnormal vessels, features that are reproduced in the well-characterized laser-induced CNV mouse model.^15^ We previously showed that in this model *Lrg1* knock-out mice developed significantly smaller lesions than those in wild type mice.^8^ In mice injected intravitreally with 15C4 immediately after lasering, fundus fluorescein angiography (FFA) was performed on day 7 (**Figure 1c**). Early phase FFA images (taken 90 s after injection of fluorescein) revealed the extent of vascularization at the site of the lesion, while late phase images (taken 7 min after injection) demonstrate fluorescein leakage, indicating hyperpermeability at the lesion. Comparison of the areas of fluorescence in the images at the two timepoints permitted quantification of the leakage ratio. Compared to control IgG injected eyes, intravitreal injection of 2.5 μg of 15C4 led to a significant reduction in vascular hyperpermeability. Intravitreal delivery of 15C4 thus leads to improved vessel function by decreasing permeability of the vessels. We also performed RNAScope^TM^ on retinal flatmounts and found *Lrg1* mRNA expression limited exclusively to endothelial cells within the lesions (**Figure 1d**), confirming that the abnormal vessels in the lesions are associated with increased expression of *Lrg1*.

### Development and analysis of Magacizumab

The high affinity of 15C4 for human LRG1, and rather lower affinity of 5.724 × 10^−7^ M for mouse LRG1 (Table 1), prompted us to map the binding epitope on human LRG1. Using a microarray of overlapping LRG1 15-mer peptides we identified the epitope in a domain located between the 6^th^ and 7^th^ leucine-rich repeats (**Figure 2a**). Alignment of the human epitope with the orthologous sequences in other mammals revealed 87% and 60% identity with cynomolgus and mouse LRG1 respectively, consistent with the relative binding affinities of 15C4 (Table 1). With a view to developing a therapeutic function-blocking antibody against LRG1 we next humanised and de-immunised 15C4, generating a hinge-stabilised IgG4 monoclonal antibody named Magacizumab. The IgG4 isotype was selected due to its low affinity for FcγRs and C1q, which would be expected to be associated with less inflammatory and complement activation properties^16^ than IgG1 or IgG2, and its intended use in the eye which is a site of immune deviation and particularly sensitive to innate immune-mediated inflammation. For development, the potential problems of Fab arm exchange and hemibody formation that may occur with therapeutic IgG4s are readily overcome with incorporation of the hinge-stabilising S228P mutation.^17–19^ Magacizumab was purified from stable-expressing CHO cells, and examined in a competition ELISA using synthetic peptides corresponding to the LRG1 epitope (**Figure 2b**). Using recombinant human LRG1 as the capture agent, the binding of Magacizumab was inhibited only by the human and primate epitope peptides, with the human peptide being an order of magnitude more effective. We also generated and tested an extended human peptide, in which the 15-mer epitope had an additional 6 amino-acids added to both N- and C-termini. However, this exhibited only slightly greater affinity for Magacizumab in the competition ELISA indicating that the major part of the epitope is captured in the original 15-mer.

**Figure 2.**
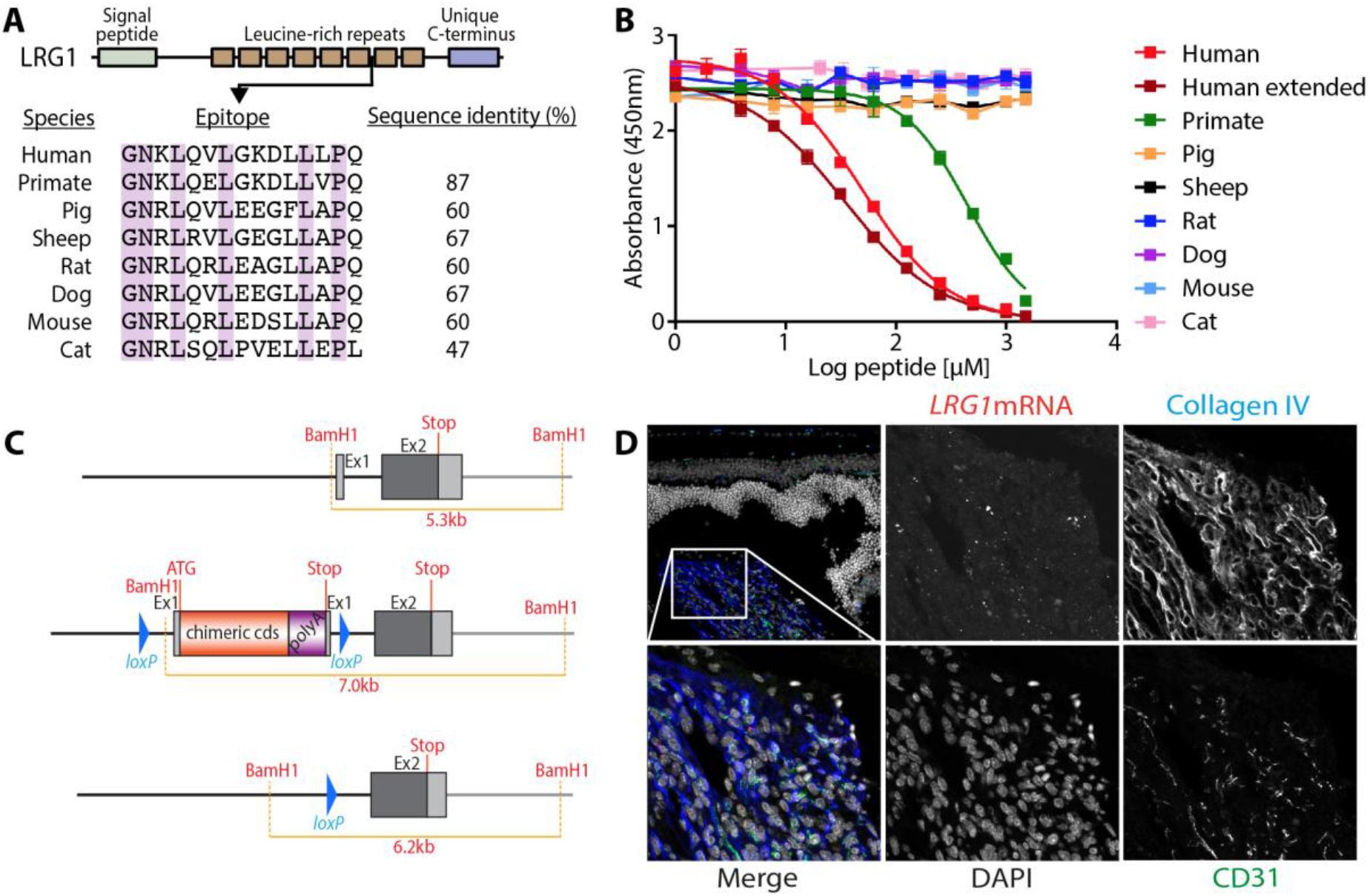
Epitope mapping and development of a human *LRG1* knock-in mouse. (**a**) Schematic representation of LRG1 showing the eight leucine-rich repeats (LRRs), with the epitope located between LRRs 6 and 7. The sequence of the human epitope is shown aligned against the orthologous sequences in seven other mammalian species, along with the sequence identity. (**b**) Competition ELISA in which Magacizumab is bound to human LRG1, with titration of the peptide epitopes of the species in A. Only the human and primate peptides were able to compete across the concentration range tested. (**c**) Schematic showing the mouse *Lrg1* gene locus, the locus following homologous recombination of the human *LRG1* targeting construct, and the locus showing loss of the chimaeric exon 1 following conditional deletion with the Cre recombinase. (**d**) Retinal sections from the human *LRG1* knock-in mouse, following laser-induced choroidal neovascularisation. The images show *LRG1* mRNA visualised with RNAscope, together with DAPI (nuclei), collagen IV and CD31. The zoomed merge image (bottom left) shows expression of human *LRG1* in the vasculature within the lesion.

SPR analysis of Magacizumab binding to human LRG1 and primate LRG1 yielded affinities of 2.81 × 10^−10^ M and 3.66 × 10^−8^ M respectively (Table 1), revealing a modest loss of affinity in the humanisation process. However, in contrast to 15C4, we were unable to detect any binding of Magacizumab to mouse LRG1, meaning that *in vivo* testing of Magacizumab would not be possible in wild type mice. To overcome this problem, we generated a human *LRG1* knock-in (h*LRG1* KI) mouse, in which a chimaeric cDNA sequence was developed utilising the mouse 5’ upstream region and coding sequence for the signal peptide, fused to a human cDNA sequence encoding the remainder of the LRG1 protein (**Figure 2c**). To demonstrate active human LRG1 expression in the knock-in mice we performed laser-induced CNV and probed the retinas using RNAScope^TM^ which revealed *LRG1* mRNA expression exclusively in the retinal lesions (**Figure 2d**). This confirms that the h*LRG1* KI mouse is a viable model for testing Magacizumab in vivo.

### Functional evaluation of Magacizumab and its Fab fragment

The capacity of Magacizumab to influence angiogenesis was validated in the foetal metatarsal explant angiogenesis assay using tissues extracted from the h*LRG1* KI mice. A reduction in angiogenesis was observed in the presence of Magacizumab (**Figure 3a**). Using human IgG4 as a control, we observed a significant and dose-dependent inhibition of both total tubule length and number of vascular junctions using Magacizumab up to a concentration of 5 μg/mL. These observations demonstrate that Magacizumab blocks angiogenesis in this assay to a similar extent as observed earlier using 15C4 with wild type mouse metatarsals. We next tested the efficacy of Magacizumab in vivo with the laser-induced model of CNV, injecting single doses of antibody from 0.01 to 1.0 μg at the time of lasering. Measurement of FFA as described earlier revealed significant inhibition of vascular leakage in 7-day lesions in the eyes of *hLRG1* KI mice injected with 1.0 μg Magacizumab (**Figure 3b**).

**Figure 3.**
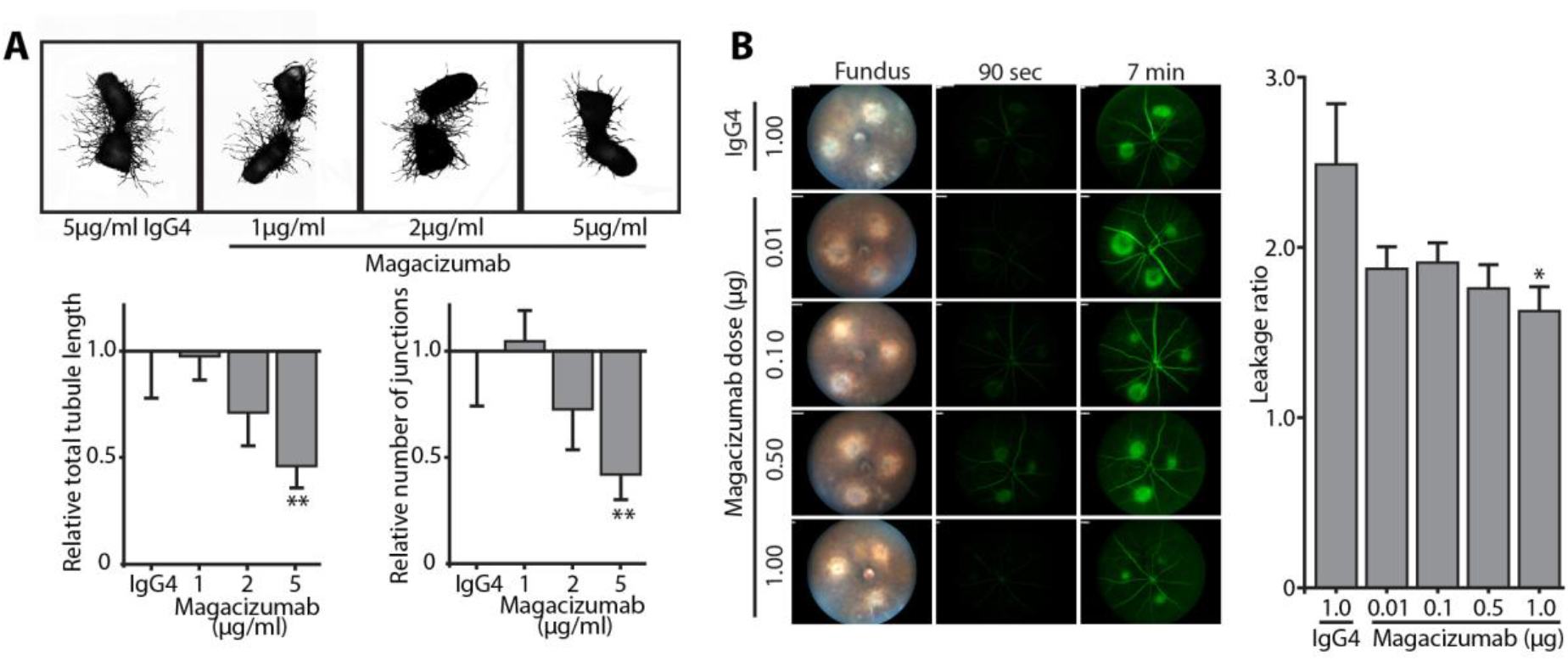
Functional evaluation of Magacizumab. (**a**) Metatarsals from human *LRG1* KI mice were treated with Magacizumab. A significant reduction in number of junctions formed and tubule length was observed in treated metatarsals when compared to IgG controls (*P* < 0.01, one-way analysis of variance). (**b**) Human *LRG1* KI mice with laser-induced choroidal neovascularisation received intravitreal injections of Magacizumab or IgG control immediately following laser burns. Fluorescein angiography was done on day 7. CNV leakage is significantly reduced in eyes treated with 1.0 μg Magacizumab compared to IgG controls (one-way ANOVA followed by Tukey’s multiple comparison test, \**P* < 0.05).

Having demonstrated the efficacy of Magacizumab in the ex vivo and in vivo models we next sought to investigate the Fab fragment of the antibody (termed MagaFab) as an alternative for therapeutic development. The treatment of ocular vascular disease has been dominated by a succession of increasingly effective anti-VEGF biologics,^20^ the first of which to be developed and most widely used is bevacizumab (Avastin), a humanised IgG1. The Fab fragment of bevacizumab, namely ranibizumab (Lucentis), was developed specifically for use in the eye and has the advantage over the parent antibody of having been through a process of affinity maturation. More recently a chimaeric molecule comprising the VEGF-binding domains of VEGFR1 and VEGFR2 fused to a human IgG1 Fc (aflibercept) has proven even more effective therapeutically, and further refinements such as brolucizumab, an anti-VEGF single chain variable fragment, are currently in clinical trials.^21^

Given that antibody Fab fragments lack potentially inflammatory Fc domains and may also be delivered at a higher molar dose due to their lower molecular weight, we prepared MagaFab from Magacizumab using papain digestion of the full-length antibody (Supplementary text Figures S3 and S4). Investigation of the affinity of MagaFab for human, primate and mouse LRG1 by SPR revealed binding kinetics and affinities similar to those of the parent antibody (Table 1). We also confirmed in ELISA that MagaFab and Magacizumab exhibited indistinguishable binding characteristics to LRG1, and also that LRG1 binding by MagaFab was unaltered upon targeted conjugation^14^ of Cy5.5 (Supplementary text Figure S5). In functional assays in h*LRG1* KI mice we observed comparable dose-dependent inhibition of vascular leakage (**Figure 4a**) to that observed for Magacizumab. Furthermore, injection of MagaFab-Cy5.5 into mouse eyes with laser-induced lesions revealed that target engagement occurs in vivo, with the Fab sequestered exclusively to the sites of neovascularisation (**Figure 4b**). This is consistent with the observed induction of *LRG1* gene expression at the lesion sites using RNAScope^TM^ (**Figure 2d**). MagaFab was also effective in the mouse foetal metatarsal assay (**Figure 4c**), using bones from h*LRG1* KI mice we observed significant inhibition of vessel tubule growth and numbers of junctions (**Figure 4d**).

**Figure 4.**
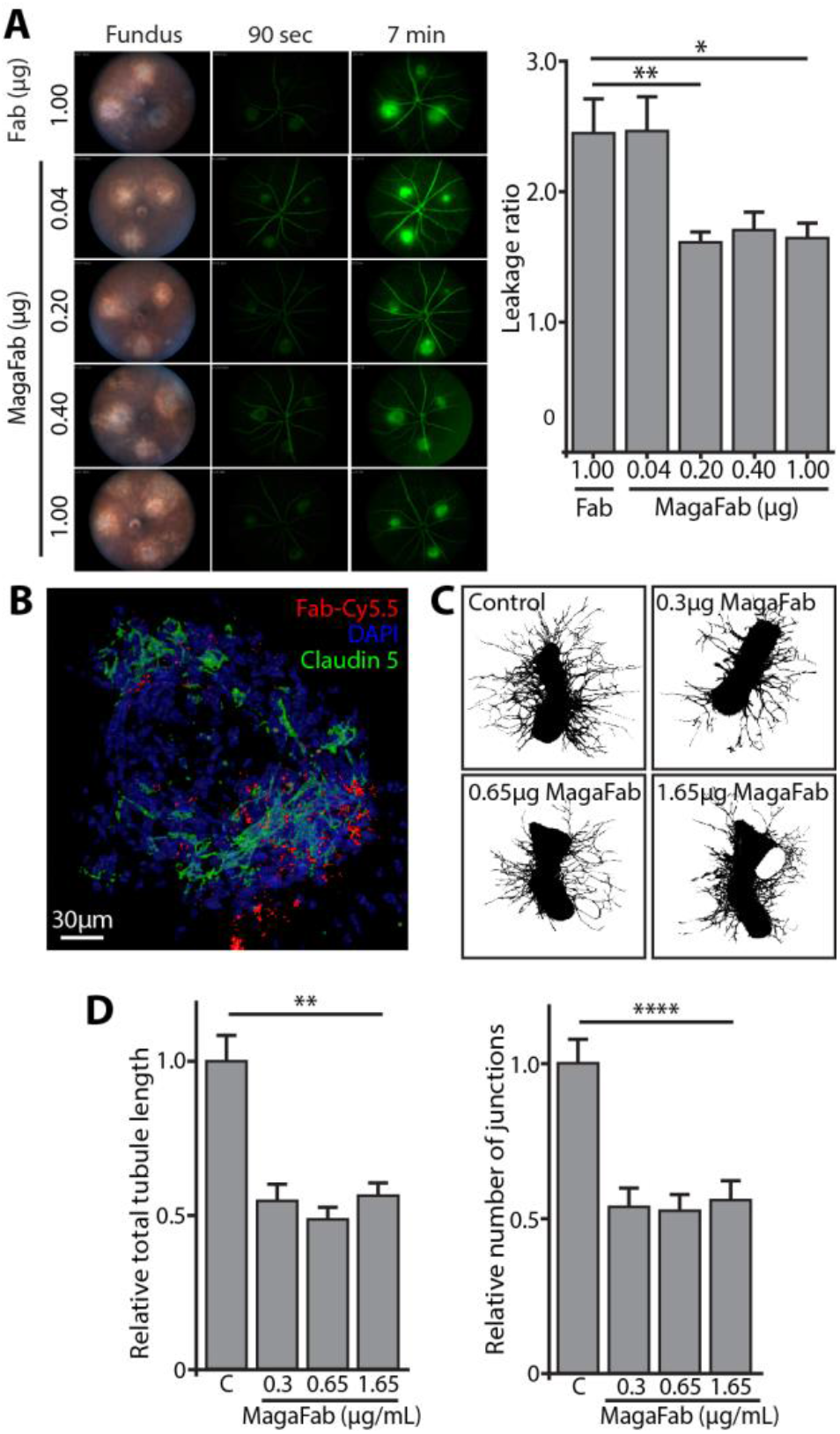
Functional evaluation of MagaFab. (**a**) Fundus fluorescein angiographic (FFA) analysis of choroidal neovascularization (CNV) lesions following an intravitreal injection of MagaFab. CNV lesions were analysed by FFA at day 7 after treatment. Images were taken 90 s after injection of dye (early FFA) and 7 min after injection of dye (late FFA) and data are presented as leakage ratio. CNV leakage is significantly reduced in eyes treated with 0.2 μg and 1.0 μg MagaFab compared to IgG controls (one-way ANOVA followed by Tukey’s multiple comparison test, \**P* < 0.05 and \*\**P* < 0.01). (**b**) Image showing the targeting of site-selectively modified MagaFab. Eyes of human *LRG1* knock-in (KI) mice were lasered and the lesions were allowed to grow for 7 days. After this time, eyes received intravitreal injections of MagaFab conjugate (Fab-Cy5.5; red). Eyes were stained for Claudin-5 (endothelial cells; green) and DAPI (nucleus; blue). (**c**) Vessel outgrowth in the metatarsal assay is attenuated by MagaFab. (**d**) Quantitation of vessel growth and branching from metatarsals treated with control Fab and MagaFab shows reduced angiogenesis in the latter. All images shown are representative and values are expressed as mean±s.e.m. of n= 3 independent experimental groups. \*\**P* < 0.001, \*\*\*\**P* < 0.0001, with all tested doses being equally effective and having identical *P* values.

## DISCUSSION

The clinical presentation of nvAMD and diabetic eye disease is characterised by a spectrum of phenotypes, and although the anti-VEGF therapeutics are generally highly effective the heterogeneity of these conditions is accompanied by a variable response to treatment.^22–24^ Attempts to generate improved therapeutics that capture poor- and non-responders can be broadly divided into two categories, one being the development of alternative and longer-acting anti-VEGFs, while other efforts have focused on alternative targets such as Ang2 and PDGF.^25^ It was a search for novel regulators of pathological angiogenesis that led us, through transcriptomic analysis of a number of rodent models of retinal vascular disease, to the discovery of LRG1.^8^ In contrast to VEGF, LRG1 is neither required for normal developmental angiogenesis, nor does it appear to have any overt homeostatic role as the null mutant mice are viable, healthy and fertile.^8^ Similar to VEGF, however, LRG1 is a secreted glycoprotein which makes it readily druggable. Moreover, as LRG1 is not normally expressed in vascular endothelial cells except in pathology, it only drives the pathogenic arm of TGFβ signaling in the disease setting. Interestingly, in evolutionary terms, one of the most closely related leucine-rich repeat proteins to LRG1 is glycoprotein A repetitions predominant (GARP or LRRC32), a cell surface protein expressed on Tregs and platelets that acts as a receptor for latent TGFβ.^26^ Monoclonal antibodies against GARP/TGFβ complexes have been shown inhibit the immunosuppressive activity of Tregs^27^ and may therefore have applications in cancer. In this study and elsewhere^27,28^ it is acknowledged that inhibition of TGFβ can be challenging because TGFβ has numerous housekeeping activities. However, targeting molecules such as LRG1 and GARP that act peripherally to the canonical TGFβ signalling pathways creates opportunities to specifically disrupt the pathogenic activities of TGFβ.

Here we generated more than 100 monoclonal antibodies against recombinant human LRG1 and screened them in a variety of functional and biophysical assays in order to identify lead antibody 15C4. As part of the analysis we performed a comparison of 15C4 with Eylea (aflibercept) in the mouse model of laser-induced CNV. Although 15C4 and Eylea apparently had comparable efficacy in our experiments, it is important to note that whilst 15C4 has high affinity for human LRG1, its affinity for mouse LRG1 is much lower. In contrast, Eylea has similar affinity for VEGF isoforms across a wide range of mammalian species including mouse.^29^ Humanisation of 15C4, to generate Magacizumab, was accompanied by a slight loss of affinity for human LRG1 and a complete loss of affinity for mouse LRG1. Ranibizumab and bevacizumab similarly have high affinity for human VEGF but do not bind to rodent VEGF.^30^ Whilst high affinity and specificity are essential in therapeutic antibodies, some loss of affinity is commonly observed following antibody humanisation.^31^ Nevertheless, this may be addressed in future work on Magacizumab with the introduction of appropriate backmutations.^32^

For clinical application in ophthalmology there are advantages in using Fab fragments rather than full-length antibodies. Accordingly, as the eye is a site of immune privilege it may be especially sensitive to innate immune responses so Fab fragments, where the Fc portion is removed, will prevent potential activation through Fc receptor engagement. In addition, given its lower molecular weight a Fab fragment may theoretically be delivered at a higher molar dose which, in spite of less favourable pharmacokinetics, will prolong the presence of an effective dose within the eye. Here we generated MagaFab and observed that it retained the binding affinity of the full-length parent antibody and was equally efficacious in experimental models. Reports that LRG1 expression is elevated in the vitreous of patients with nvAMD and diabetic eye disease,^8–10,33,35^ provide a rationale for taking MagaFab into clinical trials in these patient groups. There may also be benefits in dual targeting of LRG1 and VEGF since they represent distinct signalling pathways, either through combination therapy or the generation of a bispecific.^36^ A precedent for this strategic approach exists in faricimab, a bispecific that targets Ang2 and VEGF, and that is currently in clinical trials in diabetic macular edema.^21,37^

In summary, we have reported here the development and characterisation of a function-blocking humanised antibody against LRG1 that inhibits both pathogenic angiogenesis and vascular leakage in ex vivo and in vivo mouse models, without any detectable toxicity or inflammatory reaction in the latter. These findings validate LRG1 as a novel candidate therapeutic target in conditions characterised by retinal vascular disease such as nvAMD and diabetic eye disease.

### Study Highlights

- **What is the current knowledge on the topic?** Our current knowledge of LRG1 is that it is a secreted glycoprotein that is up-regulated in a number of different pathologies, frequently those that involve destabilisation of the vasculature, such as neovascular age-related macular disease and solid cancers. We also know that it contributes to vascular pathology by disrupting TGFβ signalling, which has led many investigators to propose that therapeutic targeting of LRG1 may be clinically beneficial.
- **What question did this study address?** Here we addressed the challenge of developing and characterising a function-blocking antibody against LRG1.
- **What does this study add to our knowledge?** We show here, not only with a mouse monoclonal but also a humanised derivative of that monoclonal, that it is possible to block the pathological activity of LRG1 in a range of *ex vivo* and *in vivo* models.
- **How might this change clinical pharmacology or translational science?** The result of our work is a novel therapeutic tool with the potential to deliver clinical benefit in a range of pathologies.

## SUPPORTING INFORMATION

Supplementary information accompanies this paper.

## AUTHOR CONTRIBUTIONS

S.E.M., J.Gr. and V.C. conceived the project, and with D.K., designed the experiments and wrote the manuscript. Experiments were performed by D.K., V.T., F.J., C.P., J.Ge., S.D., M.O’C., L.D., N.J., M.G. and A.H. Expertise and advice for the surface plasmon resonance studies were provided by F.M. and D.S., whilst S.L. and J.B. contributed expertise and advice to the experiments involving laser-induced choroidal neovascularisation. We would like to thank Stephen Perkins and Jayesh Gor (UCL) for assistance with SPR studies.

## REFERENCES

1. Chappelow, A.V. & Kaiser, P.K. Neovascular age-related macular degeneration: potential therapies. Drugs 68, 1029–1036 (2008).

2. Cohen, S.R. & Gardner, T.W. Diabetic retinopathy and diabetic macular edema. Dev. Ophthalmol. 55, 137–146 (2016).

3. Schlottmann, P.G., Alezzandrini, A.A., Zas, M., Rodriguez, F.J., Luna, J.D. & Wu, L. New treatment modalities for neovascular age-related macular degeneration. Asia Pac. J. Ophthalmol. 6, 514–519 (2017).

4. Fogli, S., Del Re, M., Rofi, E., Posarelli, C., Figus, M. & Danesi, R. Clinical pharmacology of intravitreal anti-VEGF drugs. Eye 32, 1010–1020 (2018).

5. Bahrami, B., Hong, T., Gillies, M.C. & Chang, A. Anti-VEGF therapy for diabetic eye diseases. Asia Pac. J. Ophthalmol. 6, 535–545 (2017).

6. Walshe, T.E., Saint-Geniez, M., Maharaj, A.S., Sekiyama, E., Maldonado, A.E. & D’Amore, P.A. TGF-beta is required for vascular barrier function, endothelial survival and homeostasis of the adult microvasculature. PLoS One 4, e5149 (2009).

7. Simó, R., Carrasco, E., García-Ramírez, M. & Hernández, C. Angiogenic and antiangiogenic factors in proliferative diabetic retinopathy. Curr. Diabetes Rev. 2, 71–98 (2006).

8. Wang, X. et al. LRG1 promotes angiogenesis by modulating endothelial TGF-β signalling. Nature 499, 306–311 (2013).

9. Koss, M.J. et al. Proteomics of vitreous humor of patients with exudative age-related macular degeneration. PLoS One 9, e96895 (2014).

10. Kim, T.W. et al. Proteomic analysis of the aqueous humor in age-related macular degeneration (AMD) patients. J. Proteome Res. 11, 4034–4043 (2012).

11. Zhang, J., Zhu, L., Fang, J., Ge, Z. & Li, X. LRG1 modulates epithelial-mesenchymal transition and angiogenesis in colorectal cancer via HIF-1α activation. J. Exp. Clin. Cancer Res. 35, 29 (2016).

12. Sun, D.C. et al. Leucine-rich alpha-2-glycoprotein-1, relevant with microvessel density, is an independent survival prognostic factor for stage III colorectal cancer patients: a retrospective analysis. Oncotarget 8, 66550–66558 (2017).

13. Ban, Z., He, J., Tang, Z., Zhang, L. & Xu, Z. LRG-1 enhances the migration of thyroid carcinoma cells through promotion of the epithelial-mesenchymal transition by activating MAPK/p38 signaling. Oncol. Rep. 41, 3270–3280 (2019).

14. Bahou, C. et al. Highly homogeneous antibody modification through optimisation of the synthesis and conjugation of functionalised dibromopyridazinediones. Org. Biomol. Chem. 16, 1359–1366 (2018).

15. Grossniklaus, H.E., Kang, S.J. & Berglin, L. Animal models of choroidal and retinal neovascularization. Prog. Retin. Eye Res. 29, 500–519 (2010).

16. Labrijn, A.F., Aalberse, R.C. & Schuurman, J. When binding is enough: nonactivating antibody formats. Curr. Opin. Immunol. 20, 479–485 (2008).

17. Labrijn, A.F. et al. Therapeutic IgG4 antibodies engage in Fab-arm exchange with endogenous human IgG4 in vivo. Nat. Biotechnol. 27, 767–771 (2009).

18. Silva, J.P., Vetterlein, O., Jose, J., Peters, S. & Kirby, H. The S228P mutation prevents in vivo and in vitro IgG4 Fab-arm exchange as demonstrated using a combination of novel quantitative immunoassays and physiological matrix preparation. J. Biol. Chem. 290, 5462–5469 (2015).

19. Angal, S. et al. A single amino acid substitution abolishes the heterogeneity of chimeric mouse/human (IgG4) antibody. Mol. Immunol. 30, 105–108 (1993).

20. Arnold, J.J. Age-related macular degeneration: anti-vascular endothelial growth factor treatment. BMJ. Clin. Evid. pii, 0701 (2016).

21. Al-Khersan, H., Hussain, R.M., Ciulla, T.A. & Dugel, P.U. Innovative therapies for neovascular age-related macular degeneration. Expert Opin. Pharmacother. 20, 1879–1891 (2019).

22. Bae, K., Noh, S.R., Kang, S.W., Kim, E.S. & Yu, S-Y. Angiographic subtypes of neovascular age-related macular degeneration in Korean: a new diagnostic challenge. Sci. Rep. 9, 9701 (2019).

23. Holekamp, N.M. Review of neovascular age-related macular degeneration treatment options. Am. J. Manag. Care 25, S172–S181 (2019).

24. Mansour, S.E., Browning, D.J., Wong, K., Flynn, H.W. Jr. & Bhavsar, A.R. The evolving treatment of diabetic retinopathy. Clin. Ophthalmol. 14, 653–678 (2020).

25. Ellis, M.P., Lent-Schochet, D., Lo, T. & Yiu, G. Emerging concepts in the treatment of diabetic retinopathy. Curr. Diab. Rep. 19, 137 (2019).

26. Ng, A.C. et al. Human leucine-rich repeat proteins: a genome-wide bioinformatic categorization and functional analysis in innate immunity. Proc. Natl. Acad. Sci. U. S. A. 108, 4631–4638 (2011).

27. Cuende, J. et al. Monoclonal antibodies against GARP/TGFß1 complexes inhibit the immunosuppressive activity of human regulatory T cells in vivo. Sci. Transl. Med. 7, 284 (2015).

28. Akhurst, R.J. Targeting TGF-β signaling for therapeutic gain. Cold Spring Harb. Perspect. Biol. 9, a022301 (2017).

29. Papadopoulos, N. et al. Binding and neutralization of vascular endothelial growth factor (VEGF) and related ligands by VEGF Trap, ranibizumab and bevacizumab. Angiogenesis 15, 171–185 (2012).

30. Lu, F. & Adelman, R.A. Are intravitreal bevacizumab and ranibizumab effective in a rat model of choroidal neovascularization? Graefe’s Arch. Clin. Exp. Ophthalmol. 247, 171–177 (2009).

31. Schwaigerlehner, L., Pechlaner, M., Mayrhofer, P., Oostenbrink, C. & Kunert, R. Lessons learned from merging wet lab experiments with molecular simulation to improve mAb humanization. Protein Eng. Des. Sel. 31, 257–265 (2018).

32. Margreitter, C., Mayrhofer, P., Kunert, R. & Oostenbrink, C. Antibody humanization by molecular dynamics simulations-in-silico guided selection of critical backmutations. J. Mol. Recognit. 29, 266–275 (2016).

33. Gao, B.B., Chen, X., Timothy, N., Aiello, L.P. & Feener, E.P. Characterization of the vitreous proteome in diabetes without diabetic retinopathy and diabetes with proliferative diabetic retinopathy. J. Proteome Res. 7, 2516–2525 (2008).

34. Zhang, X. et al. Leucine-rich α-2-glycoprotein predicts proliferative diabetic retinopathy in type 2 diabetes. J. Diabetes Complications 33, 651–656 (2019).

35. Chen, C. et al. Elevated plasma and vitreous levels of leucine-rich-α2-glycoprotein are associated with diabetic retinopathy progression. Acta Ophthalmologica 97, 260–264 (2019).

36. Labrijn, A.F., Janmaat, M.L., Reichert, J.M. & Parren, P.W.H.I. Bispecific antibodies: a mechanistic review of the pipeline. Nat. Rev. Drug Discov. 18, 585–608 (2019).

37. Sahni, J. et al. Simultaneous Inhibition of Angiopoietin-2 and Vascular Endothelial Growth Factor-A with Faricimab in Diabetic Macular Edema: BOULEVARD Phase 2 Randomized Trial. Ophthalmology 126, 1155–1170 (2019).

