## Supplementary data for "A Humanized Antibody against LRG1 that Inhibits Angiogenesis and Reduces Retinal Vascular Leakage"

### Appendix

### Materials and Methods

##### General experimental

All chemical reagents were purchased from Sigma Aldrich, ThermoFischer Scientific, Alfa Aesar and Acros. Compounds and solvents were used as received. Petrol refers to petroleum ether (b.p. 40–60 °C). Chemical reactions were monitored using thin layer chromatography (TLC) on pre-coated silica gel plates (254 µm) purchased from VWR. Flash column chromatography was carried out with pre-loaded GraceResolv$_{TM}$ flash cartridges on a Biotage® Isolera Spektra One flash chromatography system. $_{1}$H NMR spectra were obtained at 600 MHz or 700 MHz. $_{13}$C NMR spectra were obtained at 150 MHz or 175 MHz. All results were obtained using Bruker NMR instruments, the models are as follows: Avance III 600, Avance Neo 700. All samples were run at the default number of scans and at 21 °C. Chemical shifts (δ) for $_{1}$H NMR and $_{13}$C NMR are quoted relative to residual signals of the solvent on a parts per million (ppm) scale. Coupling constants (*J* values) are reported in Hertz (Hz) and are reported as $J_{H\text{-}H}$ couplings.

##### SDS-PAGE

Non-reducing glycine-SDS-PAGE 12% acrylamide (10% for Fab) gels were performed following standard lab procedures. A 6% stacking gel was used and a broad-range molecular weight marker (10-250 kDa) was run alongside the samples to estimate the weight of the proteins. Samples (7 µL at ~ 6 µM) were mixed with loading buffer (2 µL, composition for 6 × SDS: 1 g SDS, 1 mL glycerol, 6 mL 0.5 M Tri buffer pH 6.8, 2 mg R-250 dye) and heated at 75 °C for 5 min. The gels were run at a constant current of 30 mA for 40 minutes using 1 × SDS running buffer and stained with Coomassie blue dye.

##### UV-vis spectroscopy

Antibody concentrations, fluorophore to antibody ratios (FAR) and pyridazinedione to antibody ratios (PAR) were determined by UV-vis spectroscopy using a nanodrop ND-1000 spectrophotometer, operating at room temperature. Baseline correction was performed using sample buffer as a blank.

#### Protein LC-MS

All proteins were prepared for analysis by repeated diafiltration into ammonium acetate buffer (50 mM ammonium acetate, pH 7.0) using VivaSpin sample concentrators (GE Healthcare, 10000 MWCO) to a concentration of 6.6 µM (1.0 mg/mL). After this, the solution was diluted to 1.3 µM (0.2 mg/mL) in water and submitted to the UCL Chemistry Mass Spectrometry Facility at the Chemistry Department, UCL for analysis on the Agilent 6510 QTOF LC-MS system (Agilent, UK). 10 µL of each sample was injected into a PLRP-S, 1000 A, 8 µM, 150 mm × 2.1 mm column, which was maintained at 60 °C. The separation was achieved using mobile phase A (5% MeCN in 0.1 % formic acid) and B (95% MeCN, 5% water 0.1% formic acid) using a gradient elution. The column effluent was continuously electrosprayed into capillary ESI source of the Agilent 6510 QTOF mass spectrometer and ESI mass spectra were obtained in positive electrospray ionisation (ESI) mode using the m/z range 1,000 to 8,000 in profile mode. The raw data was converted to zero charge mass spectra using maximum entropy deconvolution algorithm using MassHunter software (version B.07.00).

#### Binding of MagaFab to LRG1 by ELISA

Binding affinity of Magacizumab Fab to LRG1 was determined by ELISA. A 96-well Maxisorp plate was coated overnight at 4 °C with LRG1 (50 µL of a 4 µg/mL solution in PBS). Next, the coating solutions were removed and each well washed with 0.1% Triton ×100 in PBS (wash buffer) three times. Then, the wells were coated with a 3% BSA solution in PBS (200 µL) for 1 h at 21 °C. After this time, the wells were emptied and washed with wash buffer 3 times. Magacizumab and Magacizumab Fab (and conjugates) were diluted in PBS yielding the following concentrations (ng/mL): 280, 140, 70, 35, 17.5, 8.75, 4.38, 2.19, 1.09, 0.547, 0.273. Wells were coated with the dilution series solutions, each in triplicate, and incubated for 2 h at 21 °C. Then, the solutions were removed, and the wells washed with wash buffer 6 times. The detection antibody; anti-human IgG (Fab-specific) HRP conjugated (1:40,000) was added and incubated for 1 h at 21 °C. Then, the solutions were removed, and the wells washed 6 times with wash buffer. Finally, equal amounts of substrate A (stabilised hydrogen peroxide) and substrate B (stabilised tetramethylbenzidine) (ELISA substrate reagent kit; R&D Systems, DY999) were premixed and added to each well (50 µL). After *ca.* 20 min the reaction was stopped by the addition of 25 µL of 2 N sulfuric acid. Absorbance was measured at 450 nm.

#### Chemical products

##### Di-*tert*-butyl-1-methylhydrazine-1,2-dicarboxylate

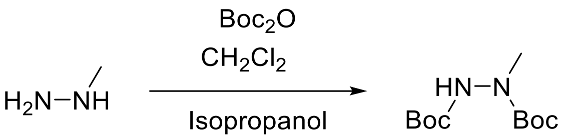


To a solution of methyl hydrazine (1.14 mL, 21.7 mmol) in IPA (16 mL), was added drop wise di-*tert*-butyl dicarbonate (11.85 g, 54.3 mmol) pre-dissolved in in $CH_2Cl_2$ (12 mL) over 30 min. The reaction was then stirred at 21 °C for 16 h. Following this, the solvents were removed *in vacuo* and the crude residue purified by flash column chromatography (0% to 20% EtOAc/petrol) to afford di-*tert*-butyl-1-methylhydrazine-1,2-dicarboxylate (4.48 g, 18.2 mmol, 84%) as a white solid **m.p.** 58–62 °C. **$^1$H NMR** (600 MHz, $CDCl_3$, rotamers) δ 6.41–6.16 (m, 1H) 3.11 (s, 3H), 1.47–1.46 (m, 18H). **$^{13}$C NMR** (150 MHz, $CDCl_3$, rotamers) δ 155.9 (C), 81.3 (C), 37.5 ($CH_3$), 28.3 ($CH_3$).

##### Di-*tert*-butyl-1-(3-(*tert*-butoxy)-3-oxopropyl)- 2-methylhydrazine-1,2-dicarboxylate

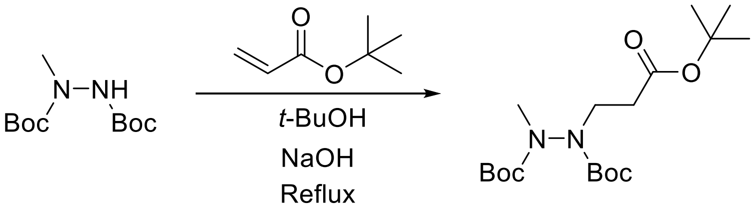


To a solution of di-*tert*-butyl 1-methylhydrazine-1,2-dicarboxylate (3.75 g, 15.2 mmol) in *tert*-butanol (25 mL), was added 0.5 mL of 2M NaOH and the reaction mixture stirred at 21 °C for 10 min. After this, *tert*-butyl acrylate (6.63 mL, 45.67 mmol) was added to the solution and the reaction mixture was heated under reflux for 72 h. The solvent was then removed *in vacuo* and the crude residue purified by flash column chromatography (0% to 20% EtOAc/petrol) to afford di-*tert*-butyl-1-(3-(*tert*-butoxy)-3-oxopropyl)-2-methylhydrazine-1,2-dicarboxylate (4.73 g, 12.6 mmol, 86%) as a clear oil. **$^1$H NMR** (700 MHz, $CDCl_3$, rotamers) δ 3.84–3.53 (m, 2H), 3.06–2.98 (m, 3H), 2.57-2.45 (m, 2H), 1.47–1.43 (m, 27H). **$^{13}$C NMR** (175 MHz, $CDCl_3$, rotamers) δ 171.1 (C), 155.5 (C), 154.5 (C), 81.1 (C), 44.7 ($CH_3$), 36.7 ($CH_2$), 34.2 ($CH_2$), 28.4 ($CH_3$).

**3-(4,5-Dibromo-2-methyl-3,6-dioxo-3,6-dihydropyridazin-1(2*H*)-yl) propanoic acid**

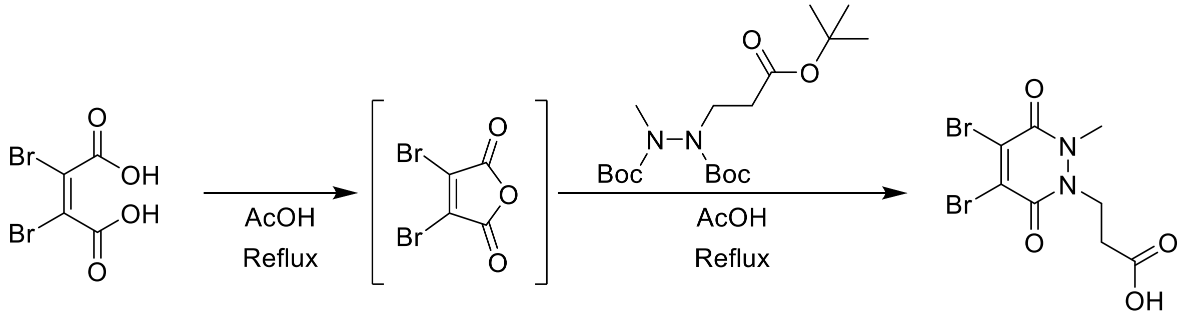


A solution of dibromomaleic acid (1.82 g, 6.68 mmol) was dissolved in AcOH (75 mL) and heated under reflux for 30 min. Di-*tert*-butyl-1-(3-(*tert*-butoxy)-3-oxopropyl)-2-methyl hydrazine-1,2-dicarboxylate (3.00 g, 8.01 mmol) was then added and the reaction heated under reflux for a further 4 h. After this time, the reaction mixture was then concentrated *in vacuo* with toluene co-evaporation (3 × 30 mL, as an azeotrope) and the crude residue purified by flash column chromatography (50% to 100% EtOAc/petrol (1% AcOH)) to afford 3-(4,5-dibromo-2-methyl-3,6-dioxo-3,6-dihydropyridazin-1(2*H*)-yl) propanoic acid (1.44 g, 4.05 mmol, 61%) as a yellow solid. **m.p.** 140–144 °C **$^{1}$H NMR** (700 MHz, MeOD) δ 4.44 (t, *J* = 7.3 Hz, 2H), 3.69 (s, 3H), 2.75 (t, *J* = 7.3 Hz, 2H). **$^{13}$C NMR** (175 MHz, MeOD) δ 173.8 (C), 154.8 (C), 154.5 (C), 136.7 (C), 136.4 (C), 44.9 ($CH_3$), 35.4 ($CH_2$), 32.6 ($CH_2$).

**2,5-Dioxopyrrolidin-1-yl 3-(4,5-dibromo-2-methyl-3,6-dioxo-3,6-dihydropyridazin-1(2*H*)-yl) propanoate**

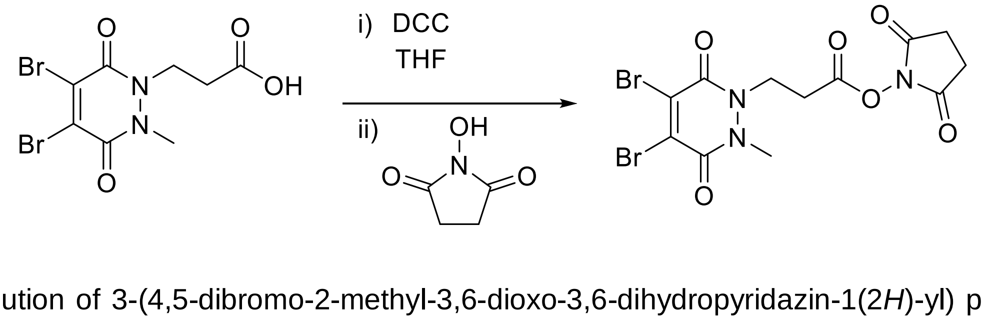


To a solution of 3-(4,5-dibromo-2-methyl-3,6-dioxo-3,6-dihydropyridazin-1(2*H*)-yl) propanoic acid (700 mg, 1.97 mmol) in THF (20 mL), pre-cooled to 0 °C, was added *N,N'*-dicyclohexylcarbodiimide (445.7 mg, 2.16 mmol). The homogenous solution was then stirred at 0 °C for 30 min. Following this, was added *N*-hydroxysuccinimide (249 mg, 2.16 mmol) and the reaction stirred at 21 °C for a further 16 h. The newly formed heterogenous mixture was then filtered and the filtrate concentrated *in vacuo*. Purification of the crude residue by flash column chromatography (30% to 100% EtOAc/petrol) afforded 2,5-dioxopyrrolidin-1-yl 3-(4,5-

dibromo-2-methyl-3,6-dioxo-3,6-dihydropyridazin-1(2*H*)-yl) propanoate (70 mg, 0.15 mmol, 8%) as a white solid. **m.p.** 100–104 °C. **$^1$H NMR** (700 MHz, $CDCl_3$) δ 4.48 (t, *J* = 6.9 Hz, 2H), 3.68 (s, 3H), 3.10 (t, *J* = 6.9 Hz, 2H), 2.85 (s, 4H). **$^{13}$C NMR** (175 MHz, $CDCl_3$) δ 168.7 (C), 166.0 (C), 153.4 (C), 153.2 (C), 136.9 (C), 135.3 (C), 43.0 ($CH_2$), 35.3 ($CH_3$), 29.1 ($CH_2$), 25.7 ($CH_2$).

**((1*R*,8*S*,9*S*)-Bicyclo[6.1.0]non-4-yn-9-yl)methyl (2-(2-(2-(3-(4,5-dibromo-2-methyl-3,6-dioxo-3,6-dihydropyridazin-1(2*H*)-yl)propanamido)ethoxy)ethoxy)ethyl) carbamate**

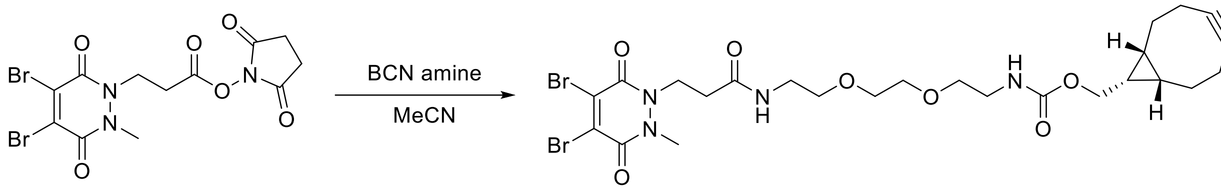


To a solution of 2,5-dioxopyrrolidin-1-yl 3-(4,5-dibromo-2-methyl-3,6-dioxo-3,6-dihydropyridazin-1(2*H*)-yl) propanoate (70 mg, 0.15 mmol) in MeCN (10 mL) was added *N*-[(1*R*,8*S*,9*S*)-bicyclo[6.1.0]non-4-yn-9-ylmethyloxycarbonyl]-1,8-diamino-3,6-dioxaoctane (50 mg, 0.15 mmol) and the reaction stirred at 21 °C for 16 h. After this time, MeCN was removed *in vacuo* and the crude residue dissolved in $CHCl_3$ (50 mL), and washed with water (2 × 30 mL) and saturated aq. $K_2CO_3$ (30 mL). The organic layer was then dried ($MgSO_4$) and concentrated *in vacuo.* Purification of the crude residue by flash column chromatography (0% to 10% MeOH/EtOAc) afforded ((1*R*,8*S*,9*S*)-Bicyclo[6.1.0]non-4-yn-9-yl)methyl (2-(2-(2-(3-(4,5-dibromo-2-methyl-3,6-dioxo-3,6-dihydropyridazin-1(2*H*)-yl)propanamido)ethoxy)ethoxy)ethyl) carbamate (73 mg, 0.11 mmol, 72%) as a yellow oil. **$^1$H NMR** (600 MHz, $CDCl_3$, rotamers) δ 7.84 (s, 0.5H), 6.34 (s, 0.5H), 5.82 (s, 0.5H), 5.29 (s, 0.5H), 4.44 (t, *J* = 6.6 Hz, 2H), 4.14–4.12 (m, 4H), 3.73–3.71 (m, 3H), 3.60–3.38 (m, 12H), 2.62 (t, *J* = 6.6 Hz, 2H), 2.27 (m, 6H), 1.61–1.57 (m, 2H), 1.39–1.24 (m, 2H), 0.96–0.94 (m, 2H). **$^{13}$C NMR** (150 MHz, $CDCl_3$, rotamers) δ 169.1 (C), 157.0 (C), 153.1 (C), 153.0 (C), 136.6 (C), 135.5 (C), 99.0 (C), 70.4 ($CH_2$), 70.3 ($CH_2$), 69.7 ($CH_2$), 63.0 ($CH_2$), 44.6 ($CH_2$), 40.8 ($CH_2$), 39.5 ($CH_2$), 35.1 ($CH_3$), 34.1 ($CH_2$), 29.3 ($CH_2$), 29.2 ($CH_2$), 21.6 ($CH_2$), 20.2 ($CH_2$), 17.9 (CH), 14.3 (CH).

### Figures

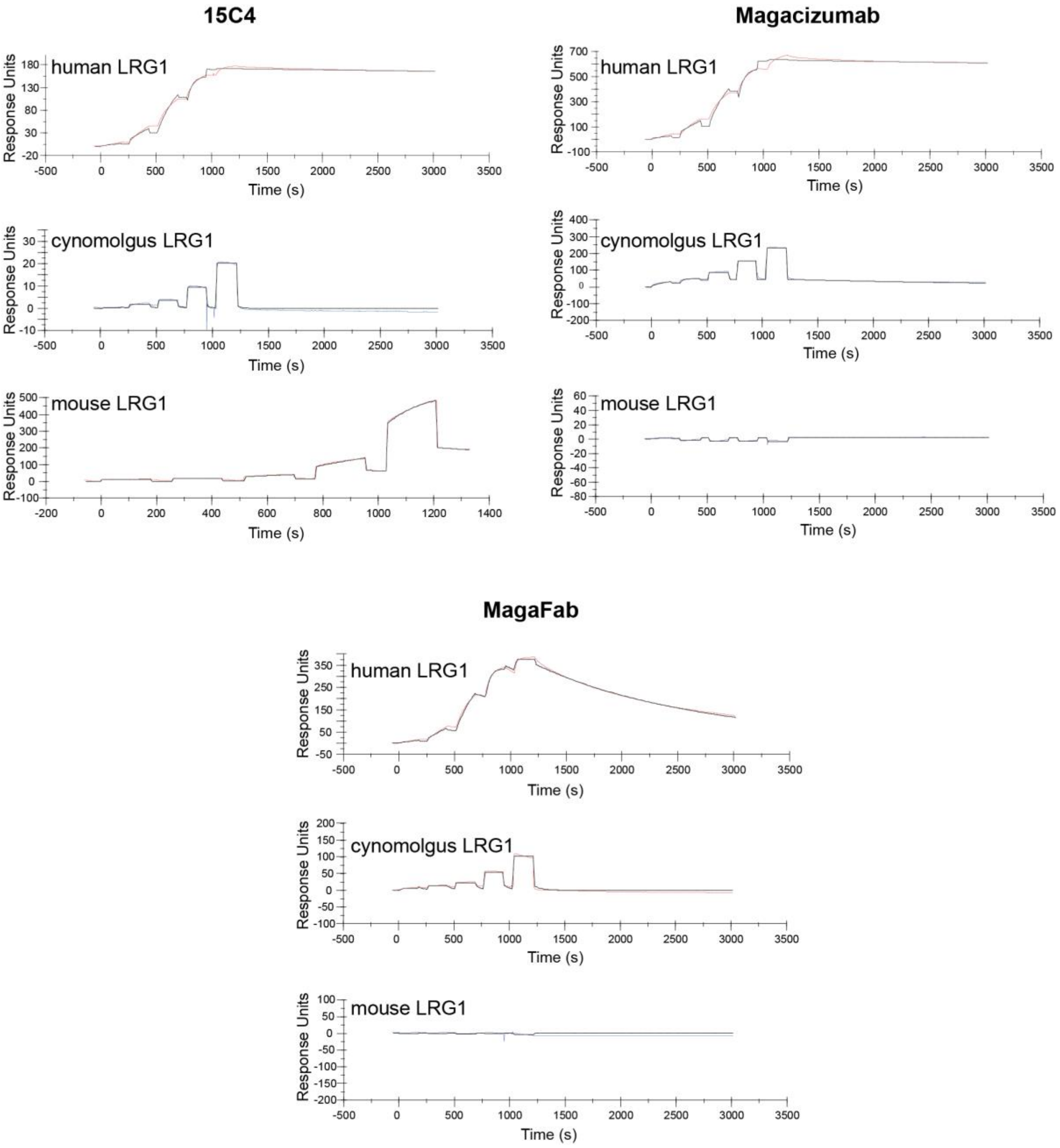


**Appendix Figure 1**

**SPR analysis of antibodies to LRG1.**

The sensorgrams show binding characteristics of 15C4, Magacizumab and MagaFab for recombinant human, cynomolgus and mouse LRG1.

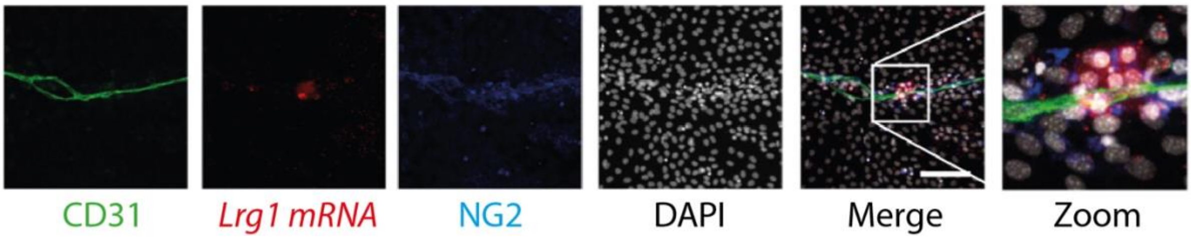


**Appendix Figure 2**

***Lrg1* expression alongside microvessels in the foetal metatarsal**.

The images show a blood vessel (CD31) growing within a field of cells in a mouse metatarsal explant. A cluster of cells expressing *Lrg1* mRNA (red) is observed in close proximity to the vessel, and to associated pericytes (blue).

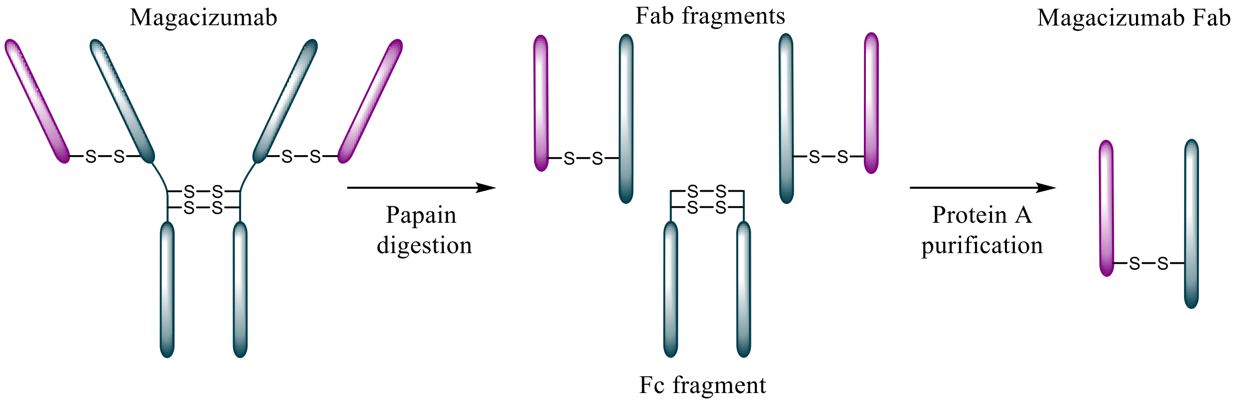


**Appendix Figure 3**

**MagaFab generation by papain digestion and protein A purification**.

Reagents and conditions: (i) Papain (1/10, papain/Magacizumab, wt/wt), Cysteine buffer pH 7.0, 37 °C, 24 h.

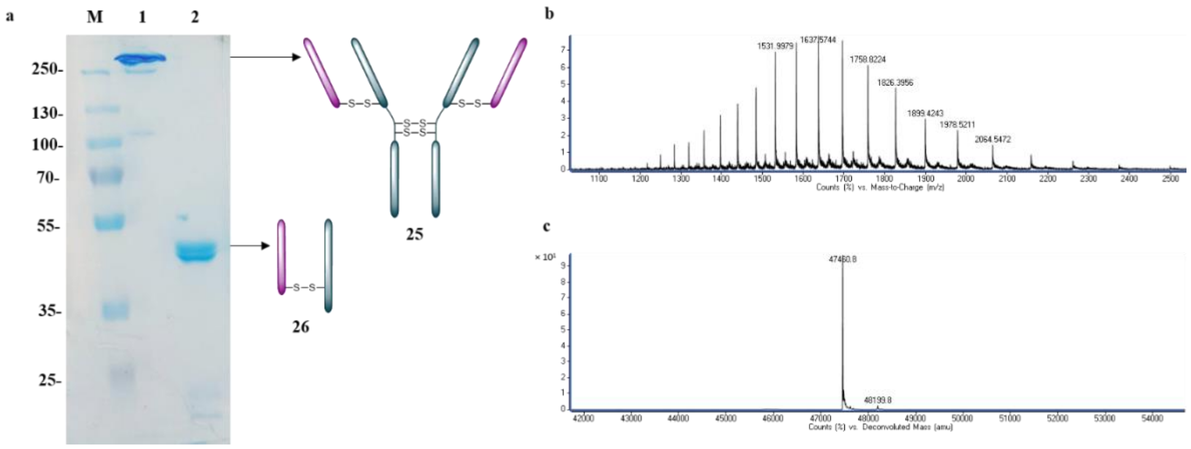


#### Appendix Figure 4

**MagaFab purity**.

A SDS-PAGE of digested Magacizumab. M – Marker; Lane 1 – Magacizumab; Lane 2 – MagaFab.

B Non-deconvoluted LCMS data for MagaFab.

C Deconvoluted LCMS data for MagaFab.

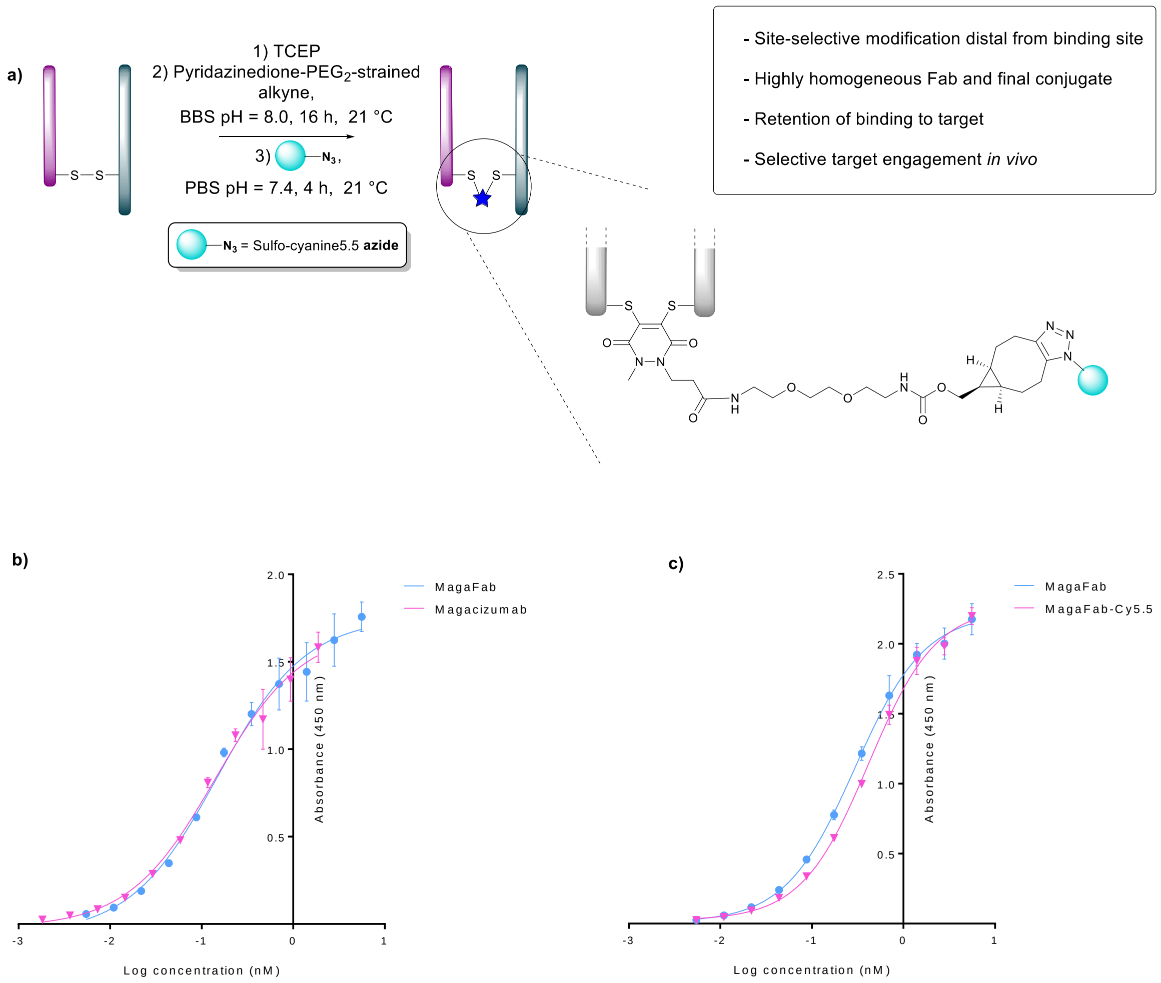


#### Appendix Figure 5

##### Synthesis of MagaFab-Cy5.5 and functional testing.

A Schematic representation of the generation of site-selectively modified Fab.

B ELISA showing that binding of MagaFab and Magacizumab to human LRG1 are equivalent.

C ELISA showing that binding of modified MagaFab Cy5.5 to human LRG1 is equivalent to that of MagaFab.
